# Rewetting of Three Drained Peatlands Drives Congruent Compositional Changes in Pro- and Eukaryotic Soil Microbiomes Through Environmental Filtering

**DOI:** 10.1101/848192

**Authors:** Micha Weil, Haitao Wang, Mia Bengtsson, Daniel Köhn, Anke Günther, Gerald Jurasinski, John Couwenberg, Wakene Negassa, Dominik Zak, Tim Urich

**Affiliations:** Institute of Microbiology, University of Greifswald, Felix-Hausdorff-Str. 8, 17487 Greifswald, Germany; Faculty of Agriculture and Environmental Sciences, University of Rostock, Justus-von-Liebig-Weg 6, 18059 Rostock, Germany; Institute of Botany and Landscape Ecology, University of Greifswald, Soldmannstraße 15, 17487 Greifswald, Germany, partner of the Greifswald Mire Center; Department of Chemical Analytics and Biogeochemistry, Leibniz-Institute of Freshwater Biology and Inland Fisheries, Müggelseedamm 301, 12587 Berlin, Germany; Department of Bioscience, Aarhus University, Vejlsøvej 25, 8600 Silkeborg, Denmark

**Author notes:** corresponding author, Institute of Microbiology, University of Greifswald, Felix-Hausdorff-Str. 8, 17487 Greifswald, Germany.

**Keywords:** peatland management, soil microbiome, methanogens, sulfate reducers, methanotrophic bacteria, greenhouse gas, methane

## Abstract

Drained peatlands are significant sources of the greenhouse gas (GHG) carbon dioxide. Rewetting is a proven strategy to protect carbon stocks; however, it can lead to increased emissions of the potent GHG methane. The response to rewetting of soil microbiomes as drivers of these processes is poorly understood, as are biotic and abiotic factors that control community composition.

We analyzed the pro- and eukaryotic microbiomes of three contrasting pairs of minerotrophic fens subject to decade-long drainage and subsequent rewetting. Also, abiotic soil properties including moisture, dissolved organic matter, methane fluxes and ecosystem respiration rates.

The composition of the microbiomes was fen-type-specific, but all rewetted sites showed higher abundance of anaerobic taxa compared to drained sites. Based on multi-variate statistics and network analyses we identified soil moisture as major driver of community composition. Furthermore, salinity drove the separation between coastal and freshwater fen communities. Methanogens were more than tenfold more abundant in rewetted than in drained sites, while their abundance was lowest in the coastal fen, likely due to competition with sulfate reducers. The microbiome compositions were reflected in methane fluxes from the sites. Our results shed light on the factors that structure fen microbiomes via environmental filtering.

## 1. Introduction

Peatlands cover 3% of the global land surface and contain about 30% of the global soil organic carbon (SOC) pool in the form of peat [1]. During the agricultural industrialization in the 20th century, many temperate mires in Europe but also other parts of the world were drained to be used as grassland or arable farming. Globally, drained or generally disturbed peatlands cover 0.3% of the global land surface but contribute approx. 5% of all anthropogenic greenhouse gas (GHG) emissions including carbon dioxide (CO_2_), methane (CH_4_) and nitrous oxide (N_2_O). The latter holds also for Germany, where drained peatlands cover approx. 5 % of the surface [2,3]. In the German federal state of Mecklenburg-West-Pomerania, peatlands cover 13% of the land surface, with their majority being drained (approx. 90%) contributing to about 30% of the total GHG budget in the region [4,5].

Peat accumulates under water-logged conditions, when more plant biomass is formed by growth than is mineralized by the prokaryotic and eukaryotic soil (micro-)biome. Water is a key factor, limiting decomposition by cutting off the supply of oxygen and other alternative terminal electron acceptors (TEA) to mineralizing microbiota [6]. Under anoxic conditions plant litter accumulates and forms peat with a high content of soil organic matter that accumulates over millennia. Changes in ecology and hydrology of peatlands can lead to substantial changes in GHG fluxes by changing the peatland biogeochemistry [7]. If peatlands are drained, formerly anoxic soil layers become aerated and carbon is released as CO_2_ during aerobic mineralization by peat microbiota [8].

In the past three decades, several 100,000 ha of European peatlands were rewetted for environmental protection and to recover some of their ecological functions [9,10]. Rewetting is a proven strategy to protect the large SOC stocks; however, it can also lead to increased emissions of the potent GHG CH_4_ and to the release of dissolved organic matter (DOM). While effects of peatland rewetting on GHG and DOM fluxes are comparably well studied [11–15], the impact on the peat microbiota, the primary driver of GHG production and emissions, is poorly understood, since it has been not well studied in temperate fens (see an exception in [16]). The major players in soil organic matter (SOM) decomposition in peat soils are microorganisms of the bacterial, archaeal and eukaryotic (e.g. fungi) domains of life, participating in a cascade of SOM degradation steps [9,17], eventually resulting in emissions of CH_4_ and CO_2_. Hydrolytic extracellular enzymes catalyze the initial decomposition of polymeric SOM. Under anoxic conditions, this step has been considered as a major bottleneck and the one determining the rate of downstream processes [8]. Further steps in the degradation are carried out through anaerobic respiratory microorganisms (like denitrifiers, iron reducers and sulfate reducers), fermentative and methanogenic microorganisms (Dean *et al.*, 2018), while methanotrophic microorganisms constitute the biological filter for CH_4_ emissions from peat (e.g. [18]). Remarkably, many novel players in soil CH_4_ cycling have been identified in the past few years [19–22]. Thus, despite more than half a century of research on methanogens, their diversity, physiology and interactions within the microbiome are still poorly understood.

It is crucial to obtain a mechanistic understanding of microbial responses and dynamics upon rewetting to suggest robust managing practices for climate-optimal peatland protection and utilization (e.g. paludiculture, [23]). Environmental filtering, i.e. the selection of certain taxa through abiotic environmental conditions, is considered to be a major mechanism structuring communities [24]. In this context, identifying the main environmental drivers of both total microbiome composition, as well as the abundance of relevant functional groups involved in carbon cycling and methanogenesis is essential.

In this study we explore the effect of rewetting on temperate fen prokaryotic and eukaryotic microbial communities. For this purpose, three pairs of drained (dry; terms in parenthesis are used in the following to refer to these ecosystems and their states) and rewetted (wet) fens with contrasting water tables and vegetation were investigated: a brackish coastal fen (Coast), a percolation fen (Perco) and an alder carr (Alder). We hypothesize that despite the obvious structural and geo genetic differences, congruent effects of rewetting will be detectable in the microbiomes of the three peatland types because of environmental filtering. Possible controlling factors of community composition, like DOM quantity and quality, soil moisture, soil/water salinity, were also determined to allow for analyzing the effect of rewetting on key players of SOM mineralization and CH_4_-cycling microorganisms.

A suite of cultivation-independent microbiological and geochemical methods was applied, including DOM profiling as well as qPCR of the methanogenesis key enzyme *mcrA* and 16S and 18S rRNA gene amplicon-based microbiome analysis. Our results may contribute to better understanding of how microbiomes respond to rewetting and to the relevant controlling environmental factors. Eventually. This could pave the way for microbiome-based proxies for predicting CH_4_ emissions from rewetted peatlands.

## 2. Materials and Methods

### 2.1 Sites and sampling

All sampling sites, alder carr, coastal fen and percolation fen, are located in north-eastern Germany (Figure 1). The alder carr is part of the Recknitz river valley, was drained at least from 1786 on and has been used as a managed forest. The rewetted sub-site (Alder_wet_) (54°13’N 12°49’E) was unintentionally flooded in 1999 due to a blocked drain and has remained wet since then. In the drained sub-site (Alder_dry_) (54°13’N 12°51’E) long-term deep drainage has caused subsidence of the soil surface of about 1 m.

**Figure 1:**
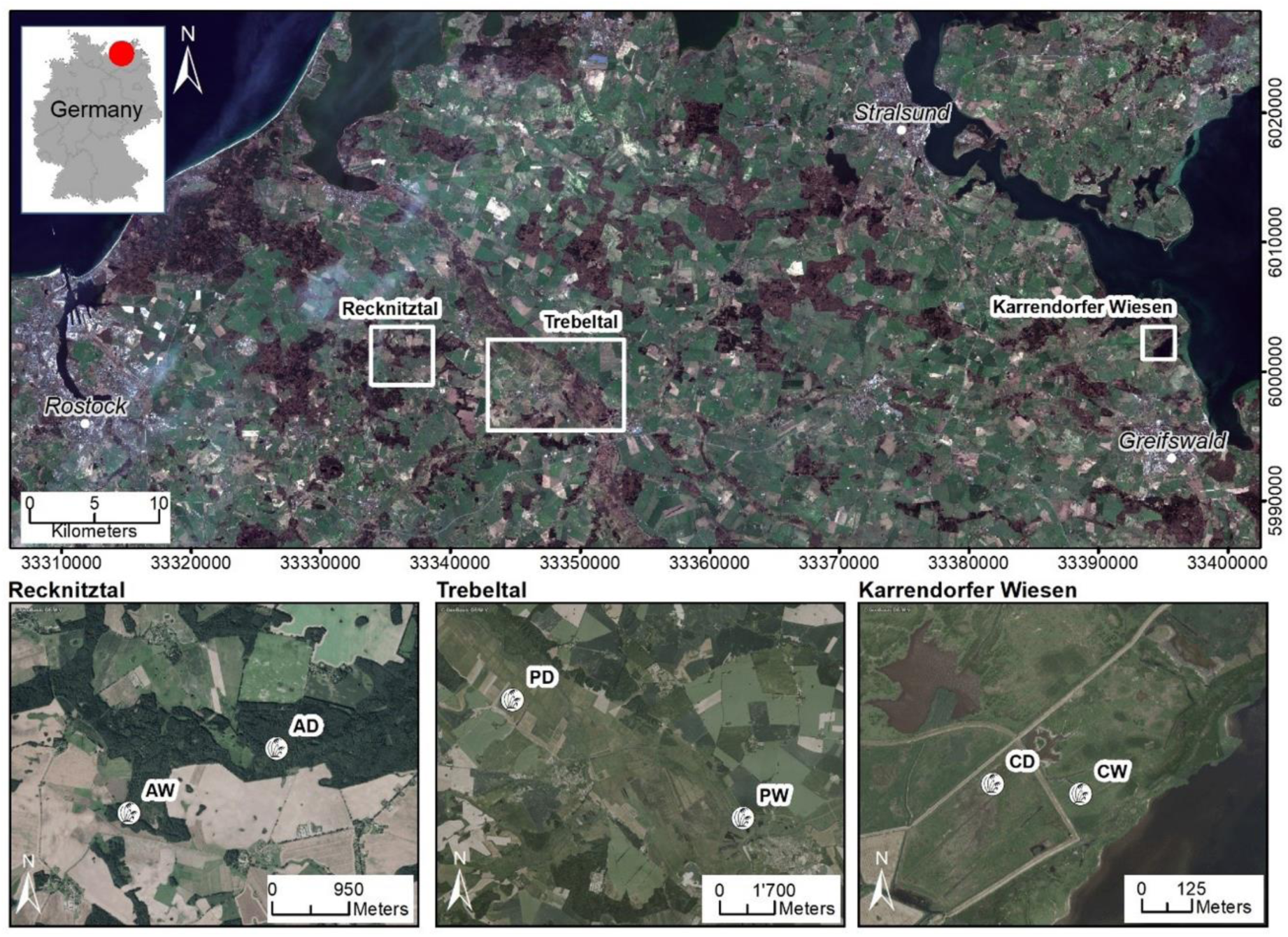
Overview of sampling sites. A: alder carr; P: percolation fen; C: coastal fen; D: drained; W: rewetted.

The coastal fen, located on a peninsula, was naturally episodically flooded by brackish seawater coming from the Bay of Greifswald of the Baltic Sea, typically during winter storm surges. The entire area was diked in 1850 to allow for pasture and other agricultural use. In 1993, the dykes around the rewetted sub site (Coast_wet_) (54°16’N 13°39’E) were removed and, subsequently, the site was episodically flooded again. The drained sub-site (Coast_dry_) (54°16’N 13°39’E) in contrast remains behind the dyke and, thus, cut off from the seawater influence. Both, Coast_wet_ and Coast_dry_ are used as cattle pastures.

The percolation fens, representing the third type of peatlands, are located in the catchment areas of the rivers Trebel and Recknitz. Both sites were deeply drained for intensive land-use and used as deep-drained grassland in the 20^th^ century. In 1998, the water table at the rewetted sub-site (Perco_wet_) (54°10’N 12°74’E) in the Trebel valley was raised again to 5-10 cm above ground level as part of a large rewetting project (EU LIFE project) (Ministerium für Bau, Landesentwicklung und Umwelt, M-V 1998). In contrast, the water table of the still drained sub-site (Perco_dry_) (54°13’N 12°63’E) in the Recknitz valley remains several decimeters below the surface and is mown 1-2 times per year.

At all sites, peat samples were taken in April 2017 as triplicate cores at 5-10 cm, 15-20 cm and 25-30 cm depths (Figure S4) using a gouge. Each sample was homogenized and stored on ice in 50-ml reaction tubes over night until further processing.

### 2.2 Soil physico-chemical properties

Soil moisture was measured gravimetrically by drying 2-3 g of soil over night at 90°C until mass constancy. Soil moisture is expressed as the percentage of lost water weight to wet soil weight. The potential soil pH was measured at room temperature with a digital pH meter (pH 540 GLP, WTW, Weilheim, Germany) in 0.01 M CaCl_2_ solution with a 1: 2.5 soil to solution ratio. Concentrations of total C, N and S were measured using a CNS analyzer (Vario MICRO cube – Elementar Analysensysteme GmbH Langenselbold, Germany). DOM was measured in soil extracts derived from mixing 3 g of previously frozen soil with 30 ml 0.1 M NaCl in 50 ml reaction tubes with subsequent shaking (vortex, 180 rpm) for 30 min. Extracts were filtered through 0.45 µm (pore size) sodium acetate filters, which were prewashed with 50 µl deionized H_2_O to remove soluble acetate. The concentrations and the composition of DOM based on size categories were determined using a size-exclusion chromatography (SEC) with organic carbon and organic nitrogen detection (LC-OCD-OND analyzer, DOC-Labor Huber, Karlsruhe, Germany) [25]. The DOM was classified into three major sub-categories: (i) ‘biopolymers’, i.e. non-humic high molecular weight substances (> 10 kDa) of hydrophilic character and no unsaturated structures like polysaccharides and proteins, (ii) aromatic ‘humic or humic-like substances’ including building blocks, and (iii) ‘low molecular-weight substances’ including low molecular weight acids and low molecular weight neutral substances. Fractions were assigned based on standards of the International Humic Substances Society. Detection limits for each fraction was 0.01 mg C L^-1^. Before analysis, all samples were stored at 5°C for less than two weeks to avoid significant changes of the DOM composition [26].

Soluble reactive phosphorus was determined with the ammonium molybdate spectrometric method (DIN EN 1189 D11) using a Cary 1E Spectrophotometer (Varian). N-NH_4_^+^ and N-NO_3_^-^ were determined calorimetrically using the photometry CFA method (Skalar SAN, Skalar Analytical B.V., The Netherlands) according to the guidelines in EN ISO 11732 (DEV-E 23) and EN ISO 13395 (DEV, D 28), respectively.

### 2.3 DNA extraction and PCR/qPCR

DNA was extracted from 0.25 g soil using the DNeasy PowerSoil Kit (QIAGEN, Hilden, Germany) according to the manufacturer’s recommendation. The bead beating step was performed using a FastPrep®-24 5G instrument (MP Biomedicals, Santa Ana, USA), with intensity 5.0 for 45 s. DNA concentrations were measured with Qubit®2.0 dsDNA High Sensitivity and dsDNA Broad Range assays (Thermo Fisher, Waltham, USA). DNA size was analyzed using 1% agarose gels.

Quantitative polymerase chain reaction was used to measure the abundances of the *mcrA* gene, coding for methyl coenzyme M reductase subunit A. Quantification of *mcrA* gene copies in peat soil DNA extracts was performed on a qTOWER 2.2 (Analytic Jena, Jena, Germany). The primer pair of forward mlas-mod and reverse *mcrA*-rev [27] was used. For each reaction, 15 µl of PCR mixture contained 7.5 µl of innuMIX qPCR MasterMix SyGreen (Analytic Jena, Jena, Germany), 0.75 µl of each primer (10 pmol/µl), 5 µl of ddH2O and 1 µl of template DNA. Duplicate measurements of two concentrations (1 and 2 ng/µl) were performed for each DNA sample. Assay condition was 95°C for 5 min, 35 cycles of 95°C for 30 s, 55°C for 45 s and 72°C for 45°C, followed by melting curve analysis to confirm PCR-product specificity. *mcrA* gene copy numbers were calculated from a standard curve obtained by serially diluting a standard from 10^6^ to 10^1^ *mcrA* gene copies per µl. The standard was created with above-mentioned primers by amplifying *mcrA* genes from cow rumen fluid [19] and cloning them into the pGEM®-Teasy vector system (Promega, Mannheim, Germany). Amplicons for *mcrA* standard were generated with vector-specific primers sp6 and T7 and the resulting PCR product was cleaned with DNA purification kit (Biozym, Hessisch Oldendorf, Germany), quality-controlled by agarose gel electrophoresis and quantified with Qubit®2.0 dsDNA High Sensitivity assay. The qPCR assay had the following parameters: slope: 3.41-3.52, efficiency: 0.92-0.96, R^2^: > 0.995.

### 2.4 Amplicon sequencing and bioinformatics

16S and 18S rRNA gene amplicons were prepared and sequenced using 300 bp paired-end read Illumina MiSeq V3 sequencing by LGC Genomics (Berlin, Germany). This comprised 16S and 18S rRNA gene PCR amplification, Illumina MiSeq library preparation and sequencing using the primer pair 515YF (GTG YCA GCM GCC GCG GTA A)/B806R (GGA CTA CNV GGG TWT CTA AT) [28] for prokaryotes and 1183F (AATTTGACTCAACRCGGG)/1443R (GRGCATCACAGACCTG) [29] for eukaryotes. The sequences have been submitted to European Nucleotide Archive (ENA), Project number PRJEB35436, accession number ERP118476, project name “Changes in peatland microbiome through rewetting”.

The data were processed using R version 3.5.1 [30]. 16S rRNA and 18S rRNA gene amplicon sequences were quality filtered, de-replicated and clustered into amplicon sequence variants (ASVs) using the *dada2* package [31]. The chimeric sequences were de-novo checked and removed with *dada2*. Then the representative sequence of each ASV was assigned to taxonomy against a modified version of the SILVA SSUref_NR_128 data base, containing an updated taxonomy for *Archaea* [32], using the programs BLASTn [33] and Megan 5 [34]. ASVs that were assigned to Chloroplast or mitochondria were filtered. Due to the low number of sequences (<500) in one sample (Coast_dry_, core 2, 15-20 cm depth) for both 16S and 18S rRNA genes, this sample was discarded from further analysis. After filtering, 1,134,953 and 3,990,919 sequences, sequences were retained and were clustered into 7697 and 5548 ASVs for 16S and 18S rRNA genes, respectively. Plant sequences in 18S rRNA genes, which accounted for 288 ASVs, were filtered for downstream analysis to increase the resolution of other eukaryotic taxa.

Alpha diversity (Shannon entropy) of the microbial community was calculated using the package *phyloseq* [35]. ASV count table was then normalized using metagenomeSeq’s cumulative sum scaling (CSS) [36]. NMDS was performed to analyze the community composition using the package *vegan* [37] with defaults. Pairwise Spearman’s rank correlations were conducted among soil properties, DNA concentrations, qPCR data, NMDS axes and methanogen relative abundances using the package *Hmisc* to detect the significance of the impact of the factors on microbial communities [38]. The soil properties with significant impact were fitted into NMDS ordination using the envfit function in the *vegan* package.

The correlations were visualized by heatmap using *corrplot*. All *P*-values for multiple comparisons were adjusted by the false discovery rate (FDR) method and the null hypothesis was rejected when *P*-values were less than 0.05.

The microbial functional groups in focus, i.e. fermenters, methanogens, methanotrophs and sulfate reducers, were predicted from 16S rRNA amplicons using FAPROTAX [39]. Also, methanogens, methanotrophs and sulfate reducers were identified based on literature knowledge; e.g. ANME2-D group within *Methanosarcina* known as methanotrophic, using the methanogenic pathway in reverse, was excluded from methanogens. All plots were created using package *ggplot2* [40].

#### 2.4.1 Co-occurrence network

A co-occurrence network was constructed to explore the potential interactions between species. Plant ASVs were included in network analysis because plant residues are a major part of peat and they might be important in interactions between microbes. To eliminate the influence of rare taxa, ASVs with relative abundances lower than 0.05% were discarded. The pairwise Spearman’s rank correlations were conducted with the package *Hmisc* and all *P*-values were adjusted by the FDR. The cut-offs of adjusted *P*-value and correlation coefficient were 0.01 and 0.7, respectively. The network was visualized using package *igraph* [41]. A network showing the connections between eukaryotes and prokaryotes was sub-selected from the original network by keeping the edges connecting these two groups. ASVs that were statistically more associated to a certain fen type were defined as indicator taxa. The indicators of each group or group combination of a factor (site or water condition) were identified using *indicspecies* package.

### 2.5 Gas flux measurements

CH_4_ exchange and ecosystem respiration (CO_2_) from soil and non-woody vegetation were measured in July 2017 with opaque, height-adjustable closed chambers in non-steady-state through-flow mode [42]. Chambers are constructed of flexible polyurethane walls and are connected to circular collars (n=5 per site, 0.63 m diameter) during the measurements. Collars were installed in the soil down to 10 cm depth more than one month prior to measurements. During chamber placement, headspace CO_2_ and CH_4_ concentrations were monitored with portable analyzers (GasScouter®, Picarro and Ultraportable Gas Analyzer, Los Gatos Research) [43].

## 3. Results

### 3.1 Study sites and soil properties

Our investigations took place in three pairs of drained and rewetted temperate peatlands in Northern Germany (see Table 1, Figure1 and materials and methods for details).

**Table 1:** Sampling site characteristics.

| Abbr. | Site | State (year of rewetting) | Water table | Peat thickness | Dominant plant species |
| --- | --- | --- | --- | --- | --- |
| Alder | Alder carr | dry | -100 cm | 60 cm | Black Alder (*Alnus glutinosa*), Ground Elder (*Aegopodium podagraria*) and Common Nettle (*Urtica dioica*) |
|  |  | wet (1999) | +15 cm | > 1 m | Black Alder (*Alnus glutinosa*), Greater Pond Sedge (*Carex riparia*) |
| Coast | Coastal fen | dry | -70 cm | 70 cm | Creeping Bentgrass (*Agrostis stolonifera*) |
|  |  | wet (1993) | -5 cm | 30 cm | Creeping Bentgrass (*Agrostis stolonifera*) |
| Perco | Percolation fen | dry | -20 cm | 6 m | Creeping Buttercup (*Ranunculus repens*), Tufted Hairgrass (*Deschampsia cespitosa*) |
|  |  | wet (1998) | +10 cm | 6 m | Lesser Pond Sedge (*Carex acutiformis*) |

In general, the gravimetric water content was higher in the rewetted than in the drained sites (Table S1). However, comparing the water content mean values of each site (mean of n=9, ± standard deviation (SD)), there was only a minor difference between Perco_dry_ (74.2% ± 5.1) and Perco_wet_ (75.4% ± 5.3), because of a high water-table near the soil surface in Perco_dry_ at the time of sampling. In all of the drained sites and in Coast_wet_, the water content increased with depth. In Perco_wet_ and Alder_wet_ (78.7 ± 4.6) with water table 10 cm above ground, the water content increased slightly with depth or did not change. The lower gravimetric water contents in Coast_dry_ (45.9% ± 6.7), Coast_wet_ (53.8% ± 10.03), and Alder_dry_ (46.3% ± 7.0) corresponded with higher amounts of mineral soil compounds (sand and clay).

The sampling sites were generally acidic to sub neutral; pH values varied between fen types but increased with depth in all drained sites. Lower pH values were detected in Coast_dry_ (3.9 - 4.0) and Alder_dry_ (4.2 - 4.7), while pH values were higher in Perco_dry_ (5.0-5.2) and Coast_wet_ (4.7 - 6.1). In Alder_wet_ (5.1 - 5.1) and Perco_wet_ (5.4 – 5.4), the pH values did not change with depth (Table S1).

Salinity in the freshwater fens was generally low (0.08 - 0.49‰). The influence of the nearby brackish bay water with a salinity of 8‰ led to a higher salinity in Coastdry (1.32‰) and in Coastwet (5.2‰) (Table S1).

DOM concentrations in the soil extracts of Alder_wet_ and Coast_wet_ were higher than in the drained sites. In Perco_wet_ DOM values measured near the surface were higher than in Perco_dry_, but they showed a strong decrease with increasing depth. The distribution of separate DOM fractions, i.e. biopolymers, humic or humic-like substances and low molecular organic matter (lCOM) followed the overall pattern of total DOM (Table S1).

Total carbon (TC) and total nitrogen (TN) content of peat decreased with depth in Alder_dry_, Coast_dry_ and Coast_wet_. In contrast, TC increased with depth in Alder_wet_, Perco_dry_ and Perco_wet_. In Alder_wet_, Perco_wet_ and Perco_dry_ TN did not change with depth. Total sulfur (TS) content increased with depth in Alder_dry_, Alder_wet_ and Perco_wet_ while it decreased with depth in Coast_dry_ and Coast_wet_. In Perco_dry_, no change was observed over depth (Table S1).

### 3.2 Microbiome analysis

#### 3.2.1 Diversity of prokaryotes and eukaryotes

Alpha diversity of prokaryotes, calculated from 16S rRNA gene amplicon sequence variants (ASVs), was significantly higher in the two freshwater peatlands than in the coastal fen (Figure S1, p < 0.001). Alpha diversity was slightly higher in rewetted than in drained sites, but this was significant only in the coastal fen (p < 0.001). Alpha diversity showed a positive correlation with soil water content, TC, TN and DNA content (Figure S2).

Similarly, alpha diversity of eukaryotes in freshwater peatlands was higher than in the coastal fen (*p* = 0.11). In contrast to prokaryotes, eukaryote alpha diversity was significantly lower in Perco_wet_ than in Perco_dry_ (*p* = 0.015), while no significant difference was observed in Alder and Coast. Alpha diversity of eukaryotes decreased with depth in the rewetted sites. For both pro- and eukaryotes, alpha diversity was negatively correlated with salinity (Figure S2).

#### 3.2.2 Microbiota composition

Community composition in 16S rRNA and 18S rRNA sequencing analysis of prokaryotes and eukaryotes showed remarkably similar patterns in the non-metric multidimensional scaling (NMDS) plot (Figure 2), with a significant clustering of microbiota from the same sampling sites (PERMANOVA *p*= 0.001). While the communities of freshwater fens Alder and Perco showed some overlap, the microbiota in Coast were clearly separated along NMDS axis 1, possibly driven by the differences in abiotic factors, such as salinity, total nitrogen (TN), total carbon (TC), C/N ratio and DOM compounds (Figure 2). Nevertheless, the pro- and eukaryotic microbiota of the dry and wet sites of each fen type were consistently separated along NMDS axis 2, indicating that the water table of the sites influenced the community composition (PERMANOVA *p*= 0.001, R^2^ = 0.267). Depth showed less impact on community composition (PERMANOVA *p* = 0.161, R^2^ = 0.046), although some clustering according to sample depth was observed.

**Figure 2:**
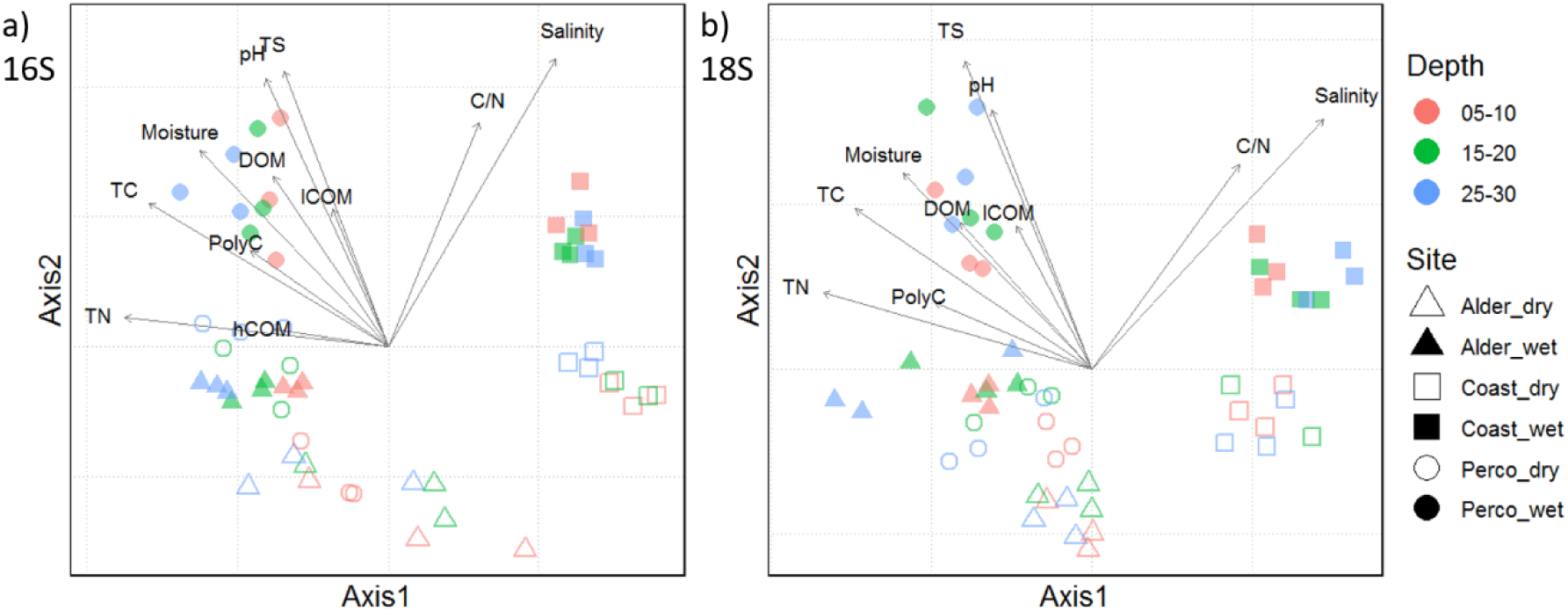
NMDS plots showing (a) prokaryotic and (b) eukaryotic community compositions. Symbols show triplicates of cores and indicate the study sites; the color indicates the depth in cm (see legend). Environmental vectors with a significance of *p* < 0.05 are shown in black. TN: total nitrogen, hCOM: humic like carbon, TC: total carbon, PolyC: Carbon polymers, DOM: dissolved organic matter, lCOM: low molecular organic matter, TS: total Sulphur, C/N: carbon/nitrogen ratio.

Pro- and eukaryotic community compositions were compared between fen types, water conditions (dry/wet) and sampling depths (Figures 3a, 5, A1). We detected a higher relative abundance of Acidobacteria in drained sites, while Betaproteobacteria, Deltaproteobacteria (mainly comprised of Myxobacteria and sulfate reducers), Chloroflexi and Bathyarchaeota showed higher values in all rewetted sites and Perco_dry_. SPAM and Latescibacteria were not detected in Coast, but they occurred in the freshwater fens, whereas Actinobacteria showed higher values in Coast than in Alder and Perco.

**Figure 3:**
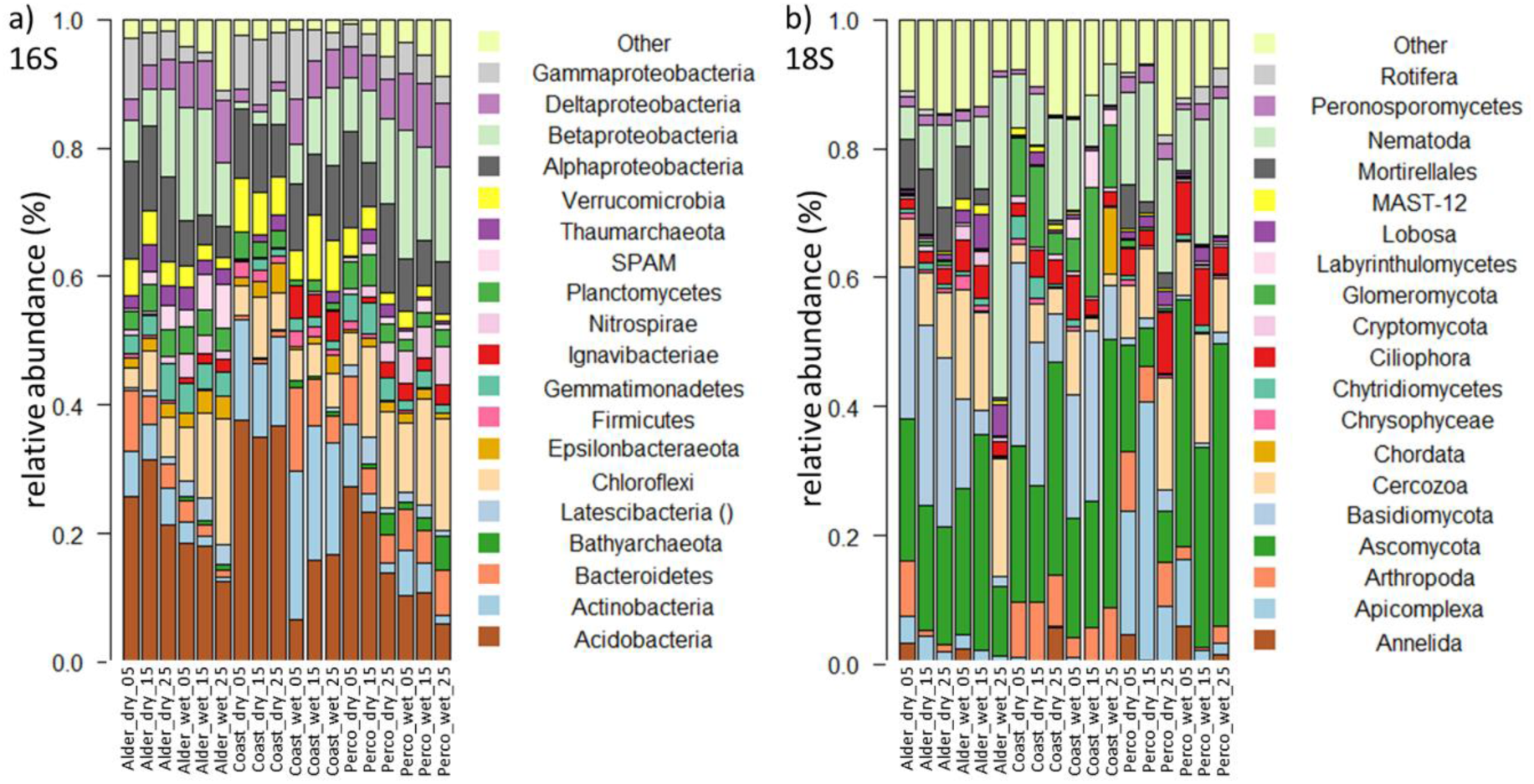
Composition of prokaryotic and eukaryotic microbial communities of drained and rewetted fens. a) Taxonomic composition of prokaryotic 16S rRNA genes displayed on phylum level, with Proteobacteria shown on class level. b) Taxonomic composition of eukaryotic 18S rRNA genes, on phylum level. Bars show triplicate of cores; phyla with < 1% abundance are displayed as “other”. The sample ID on x-axis displays fen type and depth in cm.

**Figure 4:**
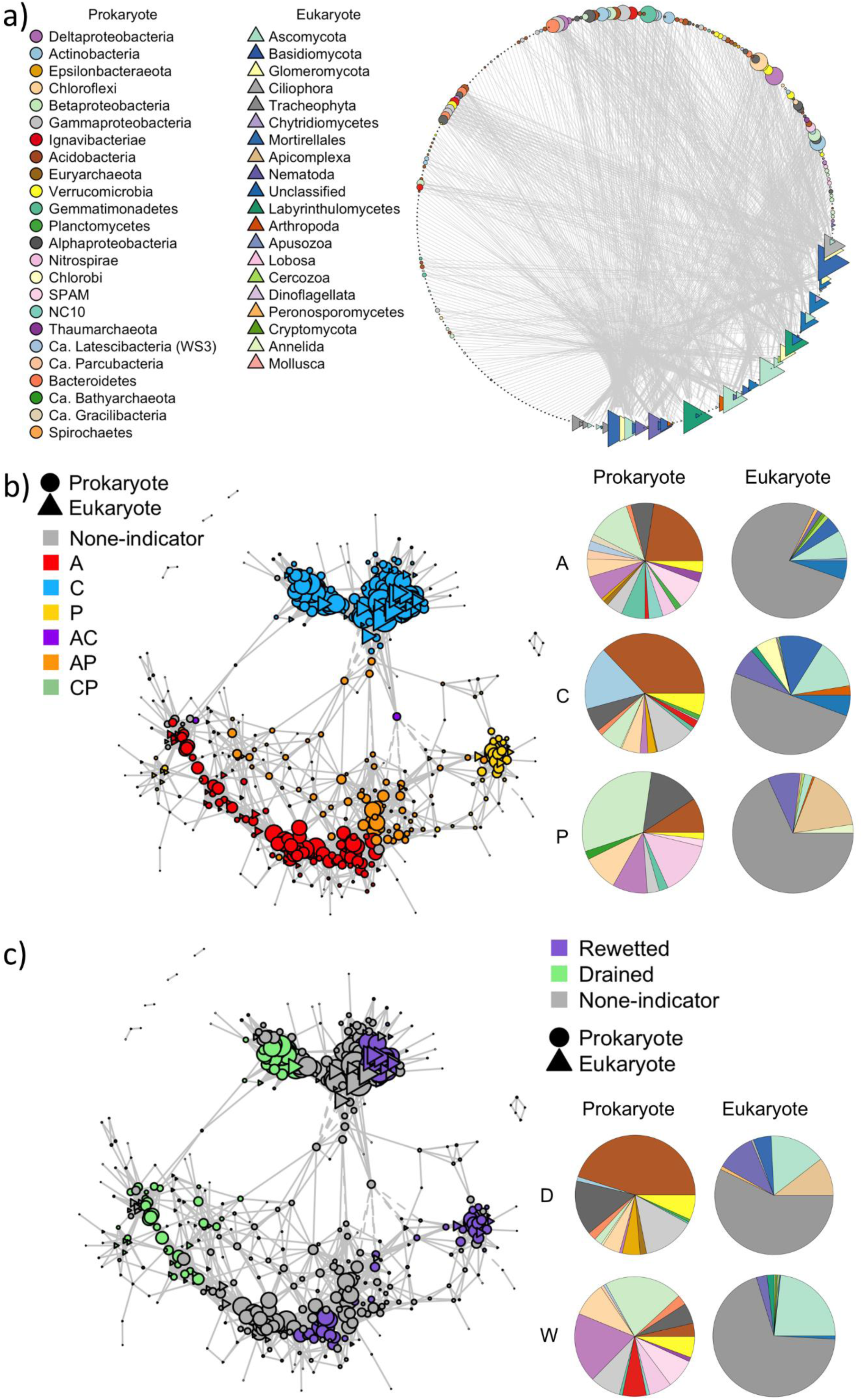
Co-occurrence networks. a) Co-occurrence network showing connections between prokaryotes and eukaryotes corresponding to the ASVs of 16S and 18S rRNA amplicon sequencing. ASVs with relative abundances lower than 0.05% were discarded. b) Co-occurrence network of ASVs associated with (A) alder carr, (C) coastal fen and (P) percolation fen, or with both alder carr and coastal fen (AC), both alder carr and percolation fen (AP) and both coastal fen and percolation fen (CP). c) Co-occurrence of ASVs associated with drained (D) and rewetted (W) sites. ASVs were associated with sites using the *indicspecies* package [44]. Solid lines indicate significant positive Spearman’s rank correlations (r > 0.7. *p* < 0.01), dashed lines indicate significant negative correlations (r < -0.7. *p* < 0.01). Nodes are sized according to the number of connected neighbors (degree) and color-coded according to site type or water condition (drained/rewetted). The relative abundance of prokaryotes and eukaryotes associated with the different site types is shown on the right-hand side of b) and c).

**Figure 5:**
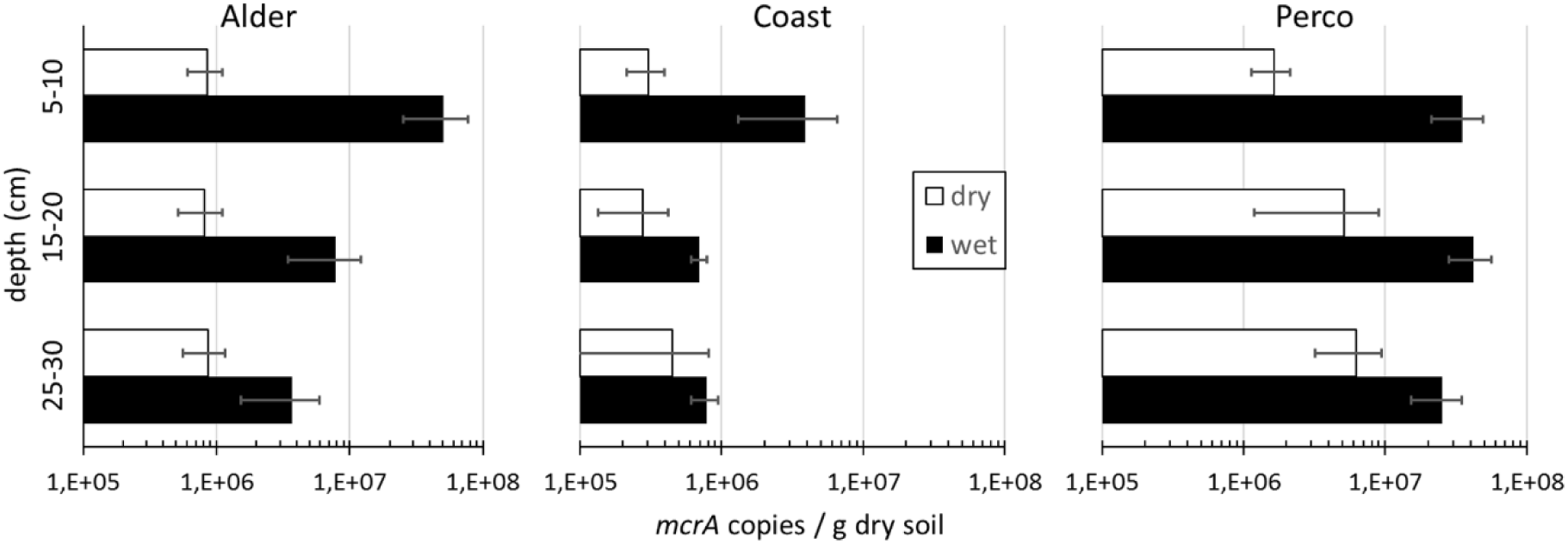
Abundance of methanogens. *mcrA* gene copy numbers per gram dry soil based on qPCR. X-axis shows mcrA copies per g dry soil, drained sites in white, rewetted sites in black bars, error bars show standard deviation of triplicate soil cores.

The eukaryotic community varied considerably between fen types and water condition (Figure 3b). Among the fungi, ASVs of Ascomycota (which contain saprophytes, yeasts and several plant pathogens) showed high relative abundance in each site, except Perco_dry_. ASV values of Basidiomycota decreased with depth in Alder and Coast, and showed higher relative abundance in these two sites than in Perco. ASV values of Glomerycota, which form arbuscular mycorrhiza with plants, comprising mostly Mortirellales were high in Alder_dry_ and Perco_dry_.

Among the protists, ASV values of Apicomplexa were high (up to 40%) in Perco_dry_, but accounted for less than 5% in other sites. Cercozoa, which contain micro-predators among other life styles, had higher values in the freshwater sites Alder and Perco. Relative abundance of Lobosa and Ciliophora, also potential micro-predators, was high in Alder_wet_ and in sites with high water content (rewetted sites and Perco_dry_) respectively.

Among the Metazoa, Nematoda showed highest relative abundance, followed by Arthropoda. The latter accounted for 5-10% of total amplicons in Coast and Perco_dry_ and the 5-10 cm layer of Alder_dry_, while their values were lower in Alder_wet_ and Perco_wet_.

#### 3.2.3 Co-occurrence network analysis

A co-occurrence network was constructed to explore the potential interactions between pro- and eukaryotic species. The network shows two distinct clusters of ASVs (Figure 4). Color coding of the nodes according to association with sites shows that ASVs within the same site were more connected than between sites (Figure 4b). However, ASVs associated with rewetted sites show little connections with those from the drained sites (Figure 4c), highlighting the significant effect of water conditions on the co-occurrence patterns. Similarly, the indicator taxa from Coast were mostly connected with each other, and indicator taxa from Perco and Alder were also distinctly connected in the network (Figure 4b). However, there were quite some shared indicator taxa from both Perco and Alder, indicating similarities between these two habitats compared with Coast, which we already detected in community composition (Figures 2 and 3). The number of a nodes adjacent edges (degree) and the number of shortest paths going through a node (closeness centrality) of the indicator taxa were higher in the wet sites than in the dry sites (Figures 4c, S4), particularly in Coast, followed by Alder, and Perco (Figures 4b, c, S4). The probability that the neighbors of a node are connected (transitivity) was higher among Coast indicators compared with those from Alder (Figure S4).

The prokaryotic indicator taxa in drained sites mainly comprised ASVs from Acidobacteria and Alpha- and Gamma-Proteobacteria, whereas the main indicator taxa of rewetted sites were Beta- and Delta-Proteobacteria. Beta- and Delta-Proteobacteria were also the main phyla in both hydrological status (Figure 3). Plants (Tracheophyta) accounted for the largest proportion of the eukaryotic indicator taxa for both water condition and site. Nematoda ASVs accounted for a larger proportion in the drained than in the rewetted sites; Ascomycota showed the reverse trend.

In the network with only connections between eukaryotes and prokaryotes (Figure 4a), eukaryotes showed higher degrees of connection compared with prokaryotes (Figure 4). Among the eukaryotes, fungi (Ascomycota, Basidiomycota and Glomeromycota) possessed the most connections with the prokaryotes.

#### 3.2.4 Functional groups

##### Methanogens

The abundance of methanogens in the samples, assessed by qPCR of the *mcrA* gene, ranged from 2.1×10^5^ to 5.1×10^7^ gene copies g^-1^ dry soil (Figure 5). The *mcrA* gene abundance was significantly higher in the three rewetted sites compared with the drained sites (Kruskal-Wallis test, H = 16.5, *p* < 0.001). The difference was largest in the top soils, where *mcrA* abundances were much higher in rewetted sites. In Alder_wet_ and Coast_wet_ methanogen abundance decreased significantly with depth, while it was rather constant in Perco_wet_. In contrast, methanogen abundance in Alder_dry_ was rather constant or increased with depth (Coast_dry_ and Perco_dry_). Positive correlations were observed between *mcrA* abundance and water, DNA and DOM content (strongest correlation with biopolymers and lCOM; Figure S2). Furthermore, TC and TN as well as pH value were positively correlated with *mcrA* abundance (Figure S2).

The relative abundance of methanogens in the prokaryotic microbiota was low. Methanogen-affiliated ASVs in the amplicon sequence datasets were on average below 0.25% of total 16S rRNA amplicons (Figure A1). In most of the drained fens, methanogen ASVs were even absent, although they were detectable by *mcrA* qPCR (Figure 5). Upper soil samples of Alder_wet_ and Coast_wet_ showed highest abundances, while methanogens were more abundant in the lower samples of Perco_wet_. Methanobacteriales and Methanosarcinales (including the families Methanosaetaceae and Methanosarcinaceae) were the dominant orders in Alder_wet_ and Coast_wet_, while methanogens were more diverse in Perco_wet_, where Methanomassiliicoccales and Methanomicrobiales were also present.

Despite apparently missing out on some methanogens in the amplicon data, the relative abundance calculated from *mcrA* gene copies per g dry soil (qPCR) shows a clear positive correlation with the relative abundance of methanogenic archaea in 16S rRNA amplicon (Spearman correlation, R= 0.578; *p* < 0.001).

##### Methanotrophs

16S rRNA genes of known methanotrophic prokaryotes as assigned via FAPROTAX analysis, were found in all sites (Figure 6), but never exceeded 1% of all prokaryotic 16S rRNA genes on average. Their relative abundance and taxon composition differed both between fen types and dry/wet. In drained fens ASVs affiliated with *Ca.* Methylacidiphilum (phylum Verrucomicrobia) were dominant across the soil profile (Figure A1). In contrast, ASVs of Methylococcaceae (class Gammaproteobacteria) were the most abundant methanotrophs in Coast_wet_ and Perco_wet_. Putative anaerobic methanotrophs affiliated with *Ca.* Methanoperedens (ANME-2d, archaea) and with *Ca*. Methylomirabilis (NC10 phylum), were the dominant methanotroph guild at the Alder_wet_ site.

**Figure 6:**
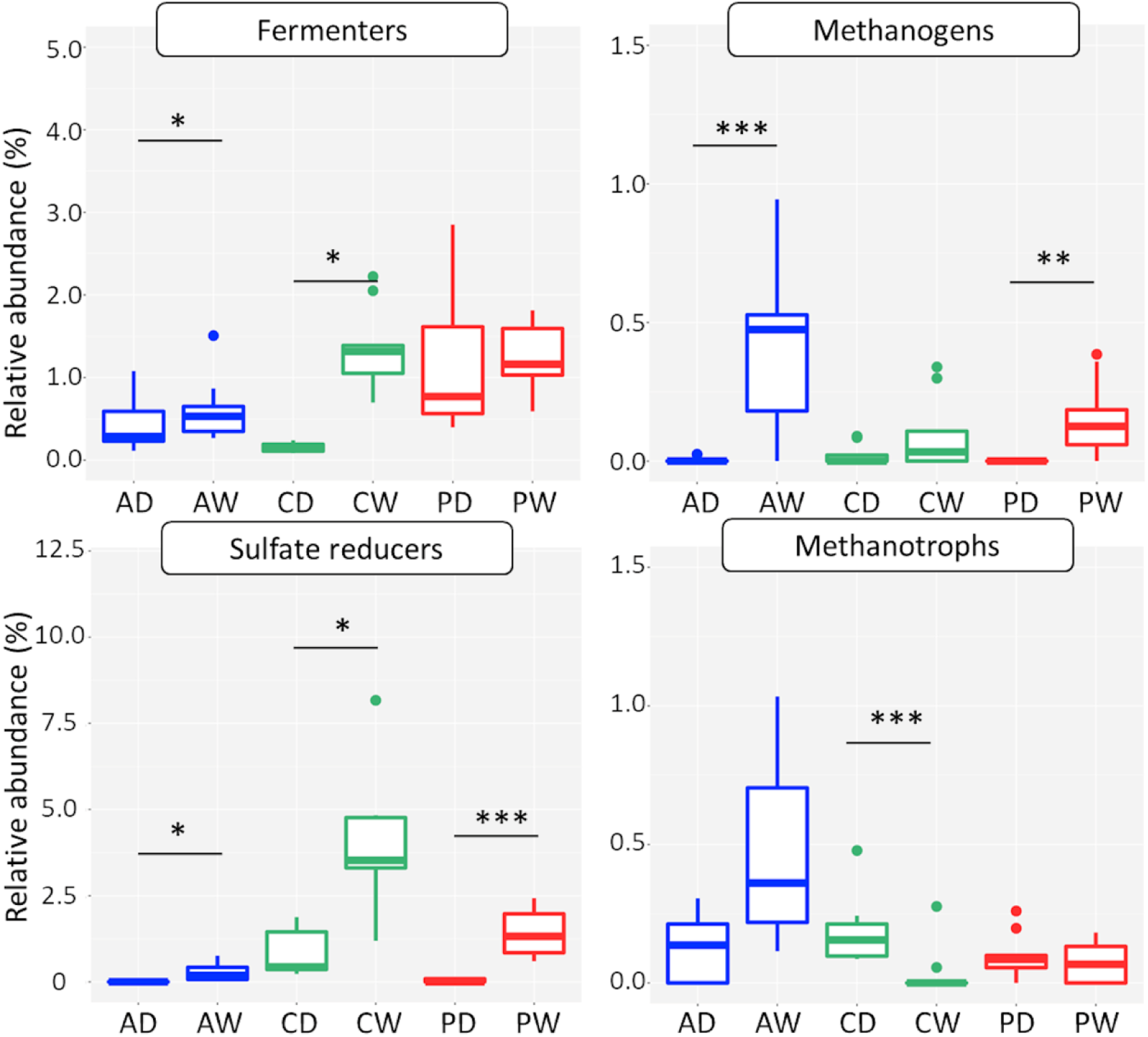
Relative abundance of functional groups in 16S rRNA gene amplicon datasets. Boxplots shows relative abundance in depth and triplicate samples. X-axes represent fen type: alder carr (A), coastal fen (C), percolation fen (P), hydrological state drained (D) and rewetted (W), depth in cm (05, 15, 25). Functional groups were assigned from 16S rRNA gene amplicon data by FAPROTAX analysis.

##### Sulfate reducers

ASVs assigned as sulfate reducers via FAPROTAX analysis were generally more abundant than methanogens and methanotrophs (between 0.1% and 5% on average, Figure 6). Like methanogens, their relative abundance was higher in rewetted sites than in drained sites. Highest relative abundance was observed in the Coast_wet_, which is exposed to recurrent influx of sea water. ASVs of Desulfobacterales were particularly abundant in this site (Figure A1). Desulfarculales and Desulfosporosinus were only detected in rewetted sites, while Desulfuromonadales occurred in all sites. Relative abundance of sulfate reducers was positively correlated to salinity, but not to total sulfur (Figure S2).

### 3.3 Greenhouse gas emissions

In all drained peatlands a net uptake of CH_4_ was measured, however, an average CH_4_ emission of 15.8 mg m^-2^ h^-1^ CH_4_ was found for Perco_wet_ with high variability (Tab. 2). Ecosystem respiration was highest in coastal peatlands. In all sites the drained peat soil emitted higher amounts of CO_2_.

## 4. Discussion

To our knowledge, this is the first comprehensive study investigating the impact of long-term rewetting (>> 10 years) on freshwater and coastal fen microbiomes. The parallel study of three different fen types allowed for the identification of overarching factors next to fen type-specific factors. The NMDS plots of both prokaryotes and eukaryotes were almost superimposable. It is remarkable that rewetting caused highly congruent effects on beta-diversity of all three domains of life and it is conceivable that water table and salinity were the main environmental filters among the physicochemical controlling factors. Rewetting resulted in more anoxic conditions in the rewetted sites compared with the drained sites. Increased anoxia is supported by several lines of evidence, including (1) a higher moisture content and (2) higher proportions of presumably fermentative microorganisms and (3) higher proportion of taxa with different types of anaerobic respiration, such as methanogenesis and sulfate reduction.

### Environmental filtering effect on pro- and eukaryotic community composition

The network analysis confirmed the conclusions we drew from the community composition: there are more taxa indicative for each of the different fen types than for water status. Consequently, site location has a more significant effect on the community assemblages. The higher degree and closeness centrality of indicators in rewetted sites indicates more connections in these sites. This might be due to that higher water flow rates in soils create greater homogeneity, and hence weak niche differentiation, contributing to stronger interactions between soil microbes. Further, it might reflect the close metabolic interactions in anaerobic microbiomes during plant biomass degradation [18,45]. Main taxa of indicator ASVs in rewetted sites were Chloroflexi and Delta- and Betaproteobacteria, which contained ASVs known from anoxic environments, also highly abundant in the wet communities (Figure 3 and Figure 4) [46]. In the drained sites, most indicator taxa were from Acidobacteria, the predominant phylum, which would correspond with the general acidic to sub-neutral site conditions. Drainage and subsequent oxidation processes in peat soils are associated with acid production and depending on the buffer capacity of soils pH may go down by one or two units compared with pre-drained conditions [47]. In contrast, rewetting with base-rich water will increase alkalinity and pH [48]. Especially the two dominant classes Acidobacteria (subdivision 1) and Solibacteres (subdivision 3) decreased in relative abundance upon rewetting (Supplementary Figure A1), which is in line with the obligatory aerobic lifestyle of most characterized member species [49]. Members of the aerobic Acidobacteria play important roles in carbohydrate degradation, but some subdivisions also contain facultative or obligate anaerobic species [49,50]. Other putative anaerobic phyla such as Ignavibacter and Bathyarchaeota occurred only in rewetted sites and deep layers of drained sites. Obviously, soil moisture acted as environmental filter leading to community compositional changes from oxygen-dependent taxa (drained) to anaerobic and fermentative taxa in the rewetted peat soils (Figure 6). A second likely environmental filter in our study is salinity. The higher connectivity and transitivity of indicators from Coast may result from environmental filtering through higher salinity in these sites, leading to a change in microbial community with increases of Actinobacteria and sulfate reducers and decrease of methanogens ([51] and see below). Remarkably, the largest proportion of eukaryotic indicator ASVs originated from Tracheophyta (plants), indicating that the interaction between plants and the soil microbiota is very important. Similar to the prokaryotes, the interaction was even stronger in rewetted sites than in drained sites. In Coast with the lowest moisture content, Tracheophyta ASVs showed lower values than in Perco and Alder. Among the animals, Nematoda indicator ASVs accounted for a larger proportion in drained sites, likely because they are dependent on oxic conditions.

In the network, we also observed that eukaryotes had higher degrees than prokaryotes. The prokaryotes mostly connected with fungi, like Ascomycota, Basidiomycota and Glomeromycota, suggesting that especially fungi may drive the composition of prokaryotic communities. We found fewer eukaryotic than prokaryotic ASVs, suggesting that one eukaryote was probably connected with multiple prokaryotes, as implied by higher degrees of connection for eukaryotes. Similarly, the richness of eukaryotes was far less than the richness of prokaryotes, which is common for soil microbiota, as shown before in e.g. [18]. Therefore, our results potentially suggest this as a universal rule of pro-eukaryote interactions.

### The influence of rewetting on methanogens

The important group of methanogenic archaea was hardly detected in the analysis of microbial networks and played no role in the determination of indicator taxa. The reason was the low relative abundance of methanogens in the amplicon datasets of the microbiota, being less than three per mil in all samples. Although their relative abundance in the microbiota was low even under rewetted conditions, their abundance per gram soil was high, especially in the rewetted sites. Wet peatlands are known to contribute a large share of the global CH_4_ emissions [1,52]. The higher *mcrA* abundance after rewetting (Figure 5) could lead to increased CH_4_ emissions from peatlands, as is the case in rice paddy fields [16,53]. Interestingly, their abundance was high in the rewetted top soils, at least in Alder_wet_ and Perco_wet_, and not in the subsoils. This is in contrast to general assumptions about their vertical distribution in the soil profile (reviewed in [6]) and corroborates recent findings from other ecosystems [54,55]. This observation makes peatland management practices such as top soil removal attractive for mitigation of CH_4_ emissions from peatlands [56,57]. The abundance of methanogens was strongly correlated with soil water content (Figure S2), which drives anoxia in the soil. However, many other factors like substrate availability of DOM, especially lCOM, C and N content of soil, degradation status of the peat and sulfate and nitrate concentration all positively correlated with methanogen abundance per gram dry soil as well (Figure S2). One significant factor was salinity, resulting from brackish water intrusions from the Baltic Sea in Coast_wet_, which led to large differences between freshwater fens (Alder and Perco) and Coast. In Coast_wet_ the abundance of methanogens per gram soil was at least 10x lower than in the freshwater sites Alder_wet_ and Perco_wet_. Thus, this environmental filtering through salinity likely leads to lower CH_4_ emissions, also shown before in several studies [46,58,59].

The strong positive correlation of *mcrA* gene abundance in qPCR with the relative abundance of methanogenic archaea in 16S rRNA gene amplicon datasets confirms the validity of the measured results. However, in most of the drained peats (Perco_dry_, most Alder_dry_ and Coast_dry_), methanogenic taxa were not present in the dataset, although *mcrA* genes were quantified by qPCR. This can be explained by the rather shallow average sequencing depth of 20,000 reads per sample, giving a taxon detection limit of about 0.05‰ in the microbiota, and thus taxa with lower abundances could not be detected. Still, the distribution of methanogenic taxa led to the assumption that substrate availability was different in the sites. While hydrogenotrophic Methanobacteriales were dominant in Coast_wet_, Coast_dry_ and Alder_wet_, Perco_wet_ showed a high variability in methanogenic groups. Here the Methanosarcinales families of Methanosaetaceae (obligate acetoclastic) and Methanosarcinaceae (flexible substrate spectrum) occurred, as did the Methanomicrobiales and the Methanomassiliicoccales (that use the H_2_ dependent methylotrophic pathway). Methanomassiliicoccales were detected in every site with Methanomassiliicoccales specific qPCR primers AS1/AS2 [60] (data not shown). Their high abundance in Perco_wet_ indicates that besides the well reported hydrogenotrophic and acetotrophic pathways [15], methylotrophic methanogenesis may contribute more to peatland CH_4_ emissions than currently assumed [19,61].

### Interactions between methanogens, methanotrophs and sulfate reducers

The second important player in peatland CH_4_ emissions is the heterogenic group of methanotrophic Bacteria and Archaea, acting as biological CH_4_ filter through aerobic or anaerobic oxidation of the produced CH_4_ (reviewed in [52,62]). Do the methanogens influence abundance of methanotrophs? The methanotrophic groups differed with site, and were generally lower abundant in the drained sites where *Ca.* Methylacidiphilum aerobic CH_4_ oxidizers were dominant. These organisms were originally isolated from acidic and volcanic environments, but have recently also been found in pH neutral environments [63,64]. They are characterized by growth on low CH_4_ concentrations, and are possibly atmospheric CH_4_ oxidizers, which would make them independent of substrate supply by methanogens. As found in other wetland studies (e.g. [65]) aerobic CH_4_ oxidizers of Methylococcaceae dominated in the wet sites. The anaerobic group *Ca.* Methylomirabilis [64], which uses oxygen from nitrite to reduce CH_4_ appeared in the sites with highest water content (Alder_wet_, Perco_dry_ and Perco_wet_). The anaerobic CH_4_ oxidizing ANME-2d (*Ca.* Methanoperedens) which reduce CH_4_ by the reverse methanogenic pathway [20] were detected in the water-covered fen Alder_wet_. Although we did not find a clear correlation between methanogen and methanotroph abundance (see [16] for similar findings), we did find a strong distinction of anaerobic and aerobic methanotrophs depending on water content and therefore oxygen availability, between sites and depths.

What is the basis of the environmental filter leading to the low methanogen abundance in the coastal fen? It is known that sulfate reducers can have an impact on methanogens, because methanogens and sulfate reducers not only share the anaerobic niche, but also compete for the same substrates, such as hydrogen [46,66,67] and acetate [68]. Due to their higher substrate affinity, sulfate reducers can outcompete methanogens, leading to lower methanogen abundance in the presence of sulfate reducers [59,68]. In fact, Desulfobacterales, known competitors for H_2_ as a substrate for methanogenesis, occurred in the coastal site Coast_wet_ with a large proportion, which was associated with a more than tenfold lower *mcrA* gene abundance than in Alder_wet_ and Perco_wet_. The influence of salinity in tidal marshes is also known to decrease CH_4_ emission compared to freshwater sites [59,69].

These inferences based on microbial analysis and soil parameters were confirmed by GHG flux measurements, conducted three months after microbial sampling. For instance, ecosystem respiration in the drained sites were significantly higher than in the wet sites, in line with the higher abundance of aerobic pro- and eukaryotes that presumably oxidize more soil organic carbon. As suggested by the higher abundance of methanogens (Table 2 and Figure 5), CH_4_ emissions were higher in wet sites compared to the drained sites. Interestingly, all drained sites as well as Coast_wet_ were CH_4_ sinks in the summer, presumably fostered by the activity of the methanotrophs. In contrast, the wet sites Alder_wet_ and Perco_wet_ with mcrA abundances < 10^7^ g^-1^ dry weight at the time of sampling acted as CH_4_ sources.

**Table 2:** Fluxes of CH_4_ and ecosystem respiration (measured as CO_2_ emissions) on all study sites compared to methanogen abundance (mcrA_DW). Negative flux values represent uptake from the atmosphere. Table shows mean values (n=5 for CH_4_ and ecosystem respiration fluxes, n=9 for mcrA_DW) and standard deviation. Significance was tested with Kruskal-Wallis test.

| Site | CH<sub>4</sub> flux (mg m<sup>-2</sup> h<sup>-1</sup>) |  | Ecosystem respiration (CO<sub>2</sub>) flux (mg m<sup>-2</sup> h<sup>-1</sup>) |  | mcrA_DW (copies g<sup>-1</sup> soil dry weight) |  |
| --- | --- | --- | --- | --- | --- | --- |
| Alder_dry | -0.08 | ± 0.07 | 493 | ± 406 | 8.5E+05 | ± 2.9E+05 |
| Alder_wet | 0.88* | ± 0.51 | 47<sup>†</sup> | ± 70 | 2.1E+07 | ± 2.6E+07 |
| Coast_dry | -0.11 | ± 0.14 | 2024 | ± 585 | 3.5E+05 | ± 2.5E+05 |
| Coast_wet | 0.00* | ± 0.01 | 1420 | ± 311 | 1.8E+06 | ± 2.1E+06 |
| Perco_dry | -0.12 | ± 0.05 | 783 | ± 152 | 4.3E+06 | ± 3.5E+06 |
| Perco_wet | 15.80* | ± 23.79 | 550<sup>†</sup> | ± 232 | 3.4E+07 | ± 1.5E+07 |

\* significantly higher than in drained site ( $p < 0.05$ )
<sup>†</sup> significantly lower than in drained site ( $p < 0.05$ )

## 5. Conclusions

In conclusion, this study provides microbiome-based insights into the environmental filtering effects of rewetting of fen ecosystems and suggests that, in terms of minimizing CH_4_ emissions, increased salinity and/or top soil removal might be promising options. Future analysis of seasonal microbiome dynamics and measurements of CH_4_ fluxes will provide evidence, whether these predictions hold true.

## Supplementary Materials

The following are available online at www.mdpi.com/xxx/s1 Figure S1: Alpha diversity of fen microbiomes, Figure S2: Heatmap showing Spearman correlation between microbial community factors and soil parameters, Figure S3 Relative abundance of Acidobacteria in 16S rRNA gene amplicon datasets, Figure S4: Node features of co-occurrence network analysis, Table S1: Biotic and abiotic soil parameters

## Author Contributions

Conceptualization, M.W. and T.U.; methodology, M.W., H.W., D.K., A.G., W.N., D.Z. and T.U.; data curation, M.W. and H.W.; formal analysis, M.W., H.W. and M.B.; funding acquisition, G.J., J.C., T.U.; investigation, M.W. and T.U.; resources, M.W., G.J., W.N., D.Z. and T.U.; visualization, M.W. and H.W.; writing—original draft, M.W., H.W., T.U.; writing—review & editing, M.W., H.W., M.B., A.G., G.J., J.C., W.N., D.Z. and T.U. All authors have read and agreed to the published version of the manuscript.

## Acknowledgment

The authors would like to thank Florian Beyer for providing the maps and arrangement of figure S4.

## Appendix A

**Figure A1:**
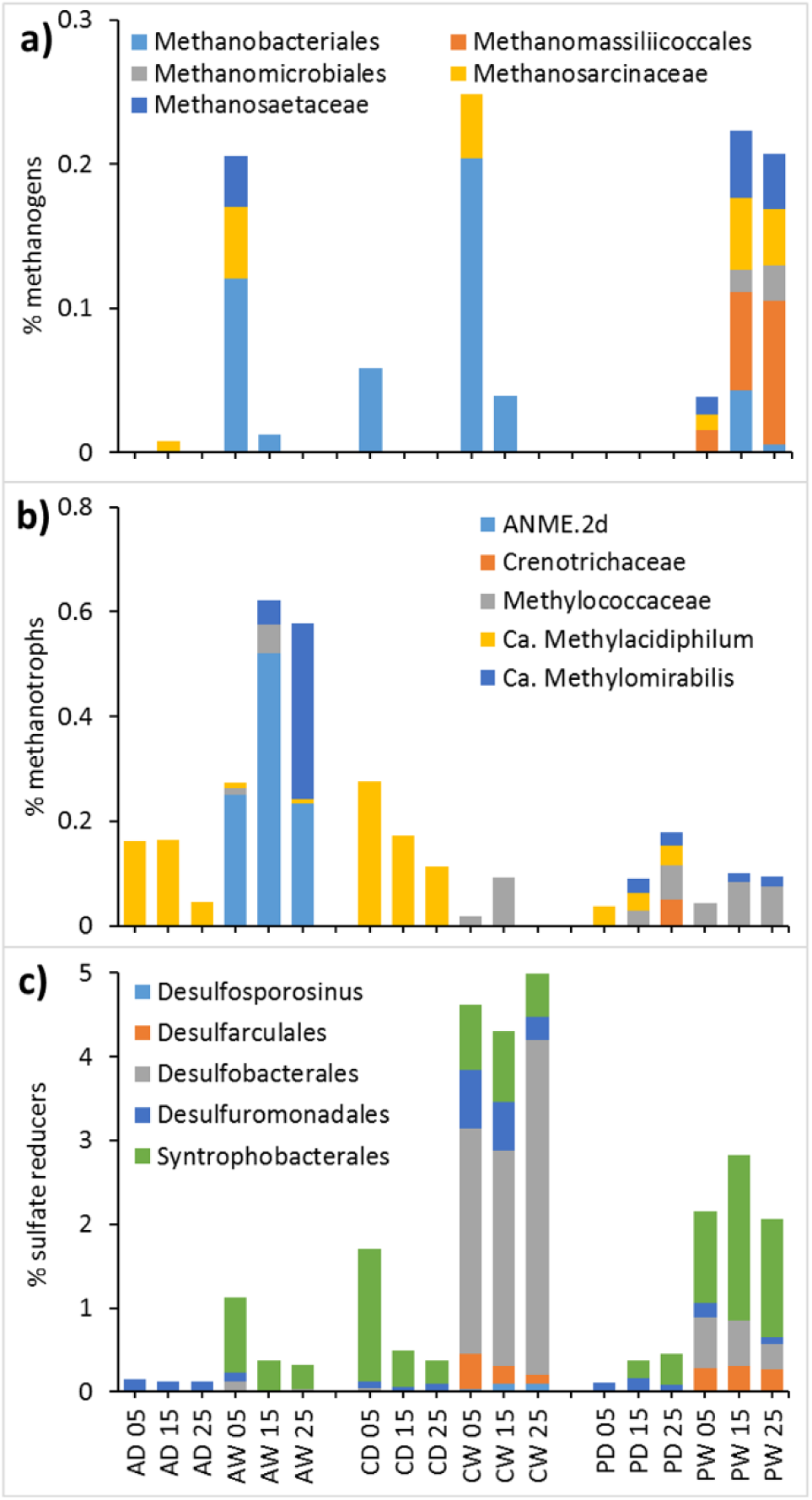
Relative abundance of functional groups in 16S rRNA gene amplicon datasets. a) Methanogens, b) methanotrophic bacteria and archaea, c) sulfate reducers. Barplots show relative abundance. X-axes represent fen type: alder carr (A), coastal fen (C), percolation fen (P), hydrological state drained (D) and rewetted (W), depth in cm (05, 15, 25). Data are shown as mean values of triplicate soil cores.

