## Supplementary figures and images for "Rewetting of Three Drained Peatlands Drives Congruent Compositional Changes in Pro- and Eukaryotic Soil Microbiomes Through Environmental Filtering"

### Figure S1

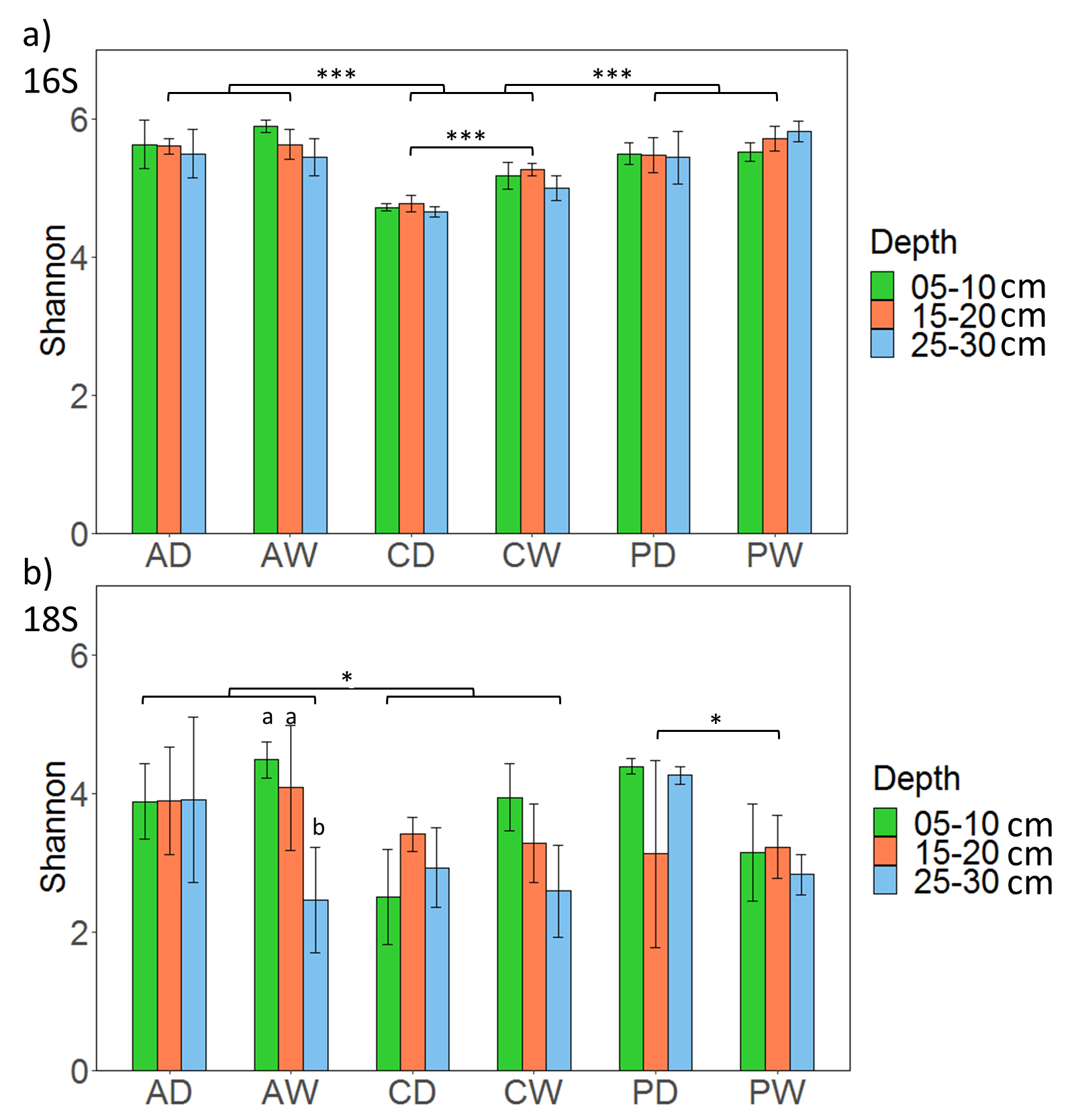

### Figure S2

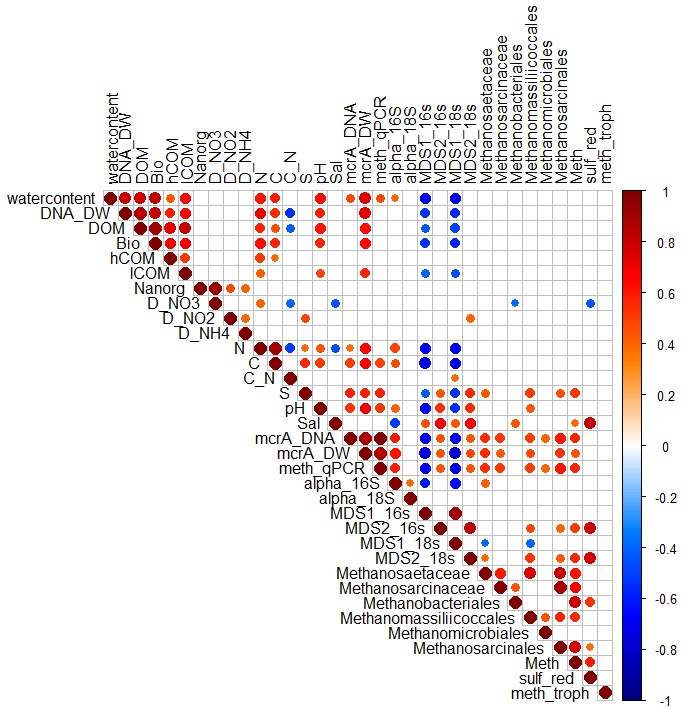

### Figure S3

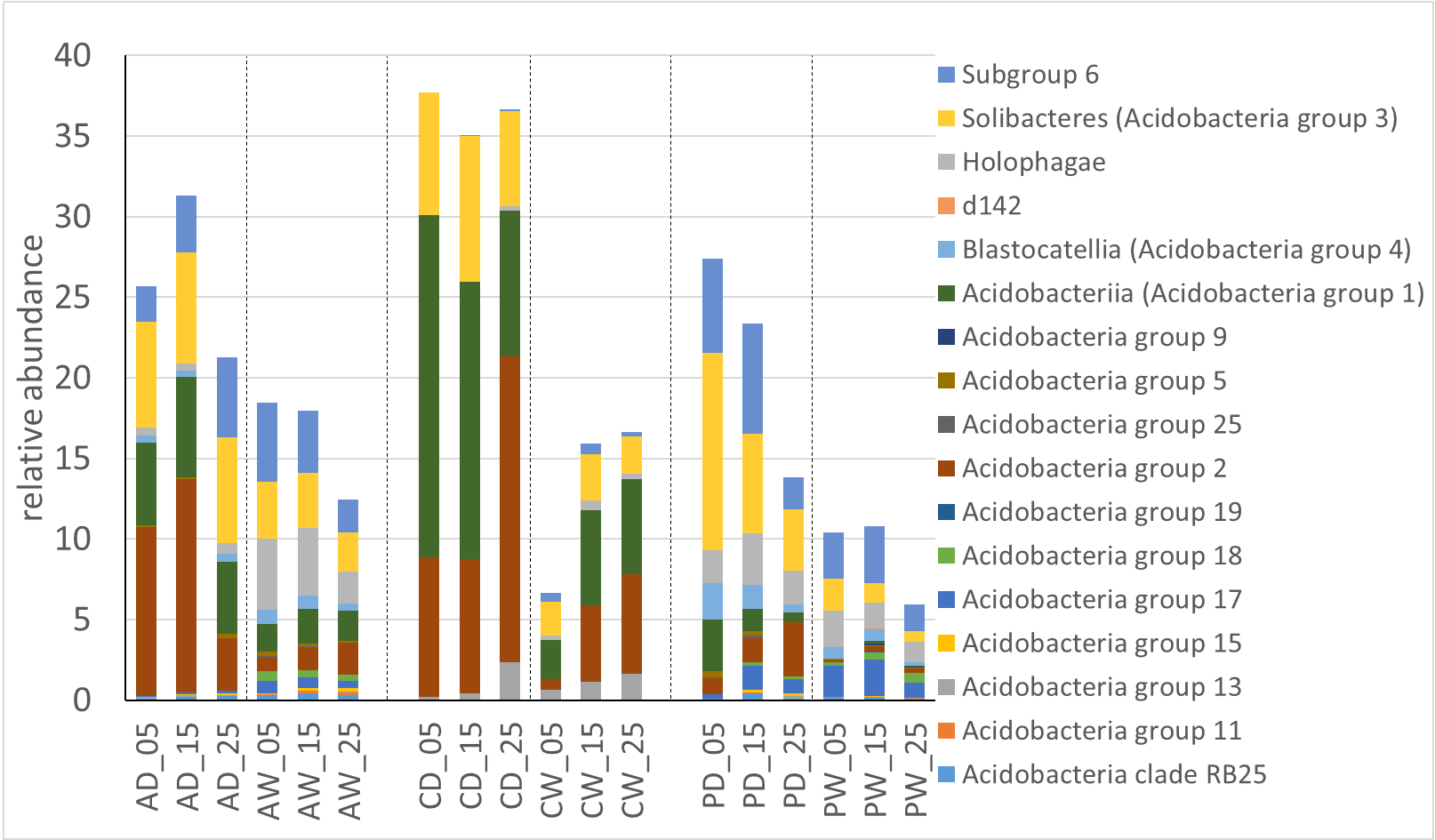

### Figure S4

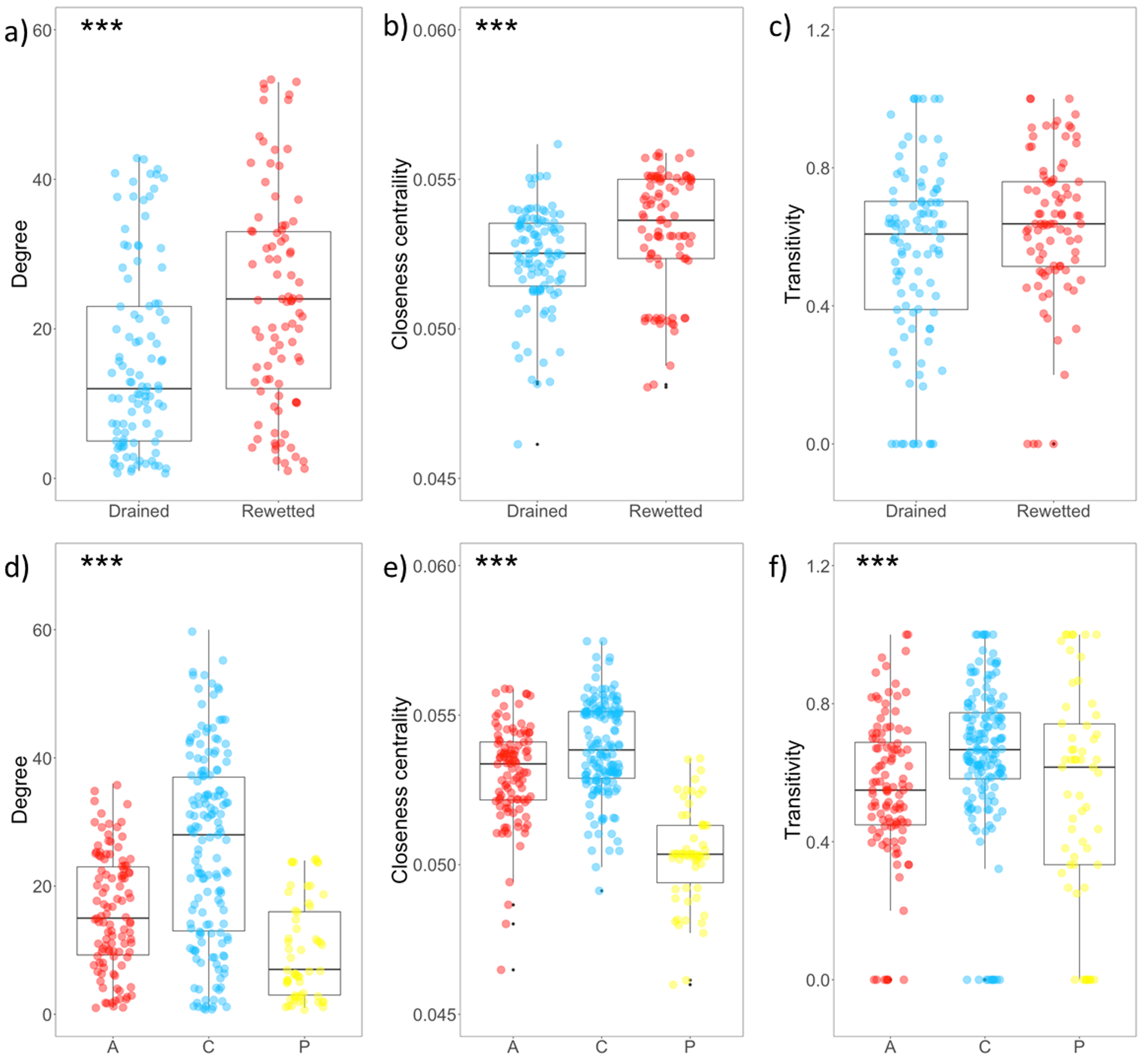
