## Supplementary material for "Rewetting of Three Drained Peatlands Drives Congruent Compositional Changes in Pro- and Eukaryotic Soil Microbiomes Through Environmental Filtering": Table S1

Table S1: Biotic and abiotic soil parameters. Abbreviations: Code: site, status, core number and depth specific code; Alder: alder swamp; Coast: coastal mire; Perco: percolation mire; dry: drained; wet: rewetted; 05, 15, 25: sampling depth in cm; moisture: gravimetric soil water content in %; DNA_DW: extracted µg DNA per gram dry weight; DOM: dissolved organic matter in µg C/g dry soil ; Bio: biopolymers in µg C/g dry soil; hCOM: high molecular carbon organic matter in µg C/g dry soil; lCOM: low molecular carbon organic matter in µg C/g dry soil; N: concentration of nitrogen in %; C: concentration of carbon in %; C_N: C/N ratio; S: concentration of sulfur in %, pHCaCl2: pH from CaCl_2_ soil extracts; Sal: salinity; mcrA_DW: mcrA gene abundance per gram dry soil.

| Code | moisture | DNA_DW | DOM | Bio | hCOM | lCOM | N | C | C_N | S | pHCaCl2 | Sal | mcrA_DW | Std_dev |
| --- | --- | --- | --- | --- | --- | --- | --- | --- | --- | --- | --- | --- | --- | --- |
| Alder_dry_1_05 | 55.4 | 32.8 | 522 | 35 | 233 | 135 | 1.27 | 16.4 | 12.9 | 0.29 | 4.20 | 0.08 | 1.21E+06 | ± 6.66E+04 |
| Alder_dry_1_15 | 48.8 | 21.2 | 291 | 25 | 138 | 85 | 0.92 | 12.9 | 14.0 | 0.24 | 4.33 | 0.08 | 1.24E+06 | ± 9.92E+04 |
| Alder_dry_1_25 | 47.8 | 33.4 | 371 | 32 | 195 | 79 | 1.98 | 29.5 | 14.9 | 0.55 | 4.72 | 0.08 | 6.67E+05 | ± 7.08E+04 |
| Alder_dry_2_05 | 42.0 | 36.9 | 396 | 24 | 215 | 75 | 1.27 | 16.4 | 12.9 | 0.29 | 4.20 | 0.08 | 7.09E+05 | ± 5.13E+03 |
| Alder_dry_2_15 | 36.0 | 25.4 | 368 | 21 | 162 | 68 | 0.92 | 12.9 | 14.0 | 0.24 | 4.33 | 0.08 | 5.41E+05 | ± 1.31E+03 |
| Alder_dry_2_25 | 37.2 | 27.7 | 535 | 41 | 247 | 148 | 1.98 | 29.5 | 14.9 | 0.55 | 4.72 | 0.08 | 6.29E+05 | ± 0.00E+00 |
| Alder_dry_3_05 | 45.9 | 21.3 | 408 | 25 | 140 | 172 | 1.27 | 16.4 | 12.9 | 0.29 | 4.20 | 0.08 | 6.56E+05 | ± 4.92E+04 |
| Alder_dry_3_15 | 45.0 | 16.6 | 180 | 14 | 81 | 47 | 0.92 | 12.9 | 14.0 | 0.24 | 4.33 | 0.08 | 6.76E+05 | ± 5.12E+04 |
| Alder_dry_3_25 | 58.3 | 51.1 | 367 | 32 | 177 | 77 | 1.98 | 29.5 | 14.9 | 0.55 | 4.72 | 0.08 | 1.30E+06 | ± 1.27E+05 |
| Alder_wet_1_05 | 89.7 | 167 | 1289 | 250 | 539 | 335 | 2.77 | 34.0 | 12.3 | 0.55 | 5.07 | 0.33 | 7.59E+07 | ± 6.28E+06 |
| Alder_wet_1_15 | 76.1 | 68.2 | 433 | 59 | 195 | 103 | 2.77 | 34.0 | 12.3 | 0.55 | 5.07 | 0.33 | 1.16E+07 | ± 2.54E+03 |
| Alder_wet_1_25 | 73.5 | 32.9 | 795 | 222 | 339 | 133 | 2.71 | 39.0 | 14.4 | 0.65 | 5.09 | 0.33 | 3.02E+06 | ± 4.30E+05 |
| Alder_wet_2_05 | 79.1 | 89.6 | 844 | 64 | 274 | 187 | 2.77 | 34.0 | 12.3 | 0.55 | 5.07 | 0.33 | 1.53E+07 | ± 7.85E+05 |
| Alder_wet_2_15 | 75.1 | 66.2 | 767 | 100 | 266 | 400 | 2.77 | 34.0 | 12.3 | 0.55 | 5.07 | 0.33 | 1.69E+06 | ± 8.75E+04 |
| Alder_wet_2_25 | 81.5 | 63.3 | 761 | 109 | 381 | 138 | 2.71 | 39.0 | 14.4 | 0.65 | 5.09 | 0.33 | 1.44E+06 | ± 2.81E+05 |
| Alder_wet_3_05 | 78.0 | 167.5 | 1025 | 178 | 555 | 138 | 2.77 | 34.0 | 12.3 | 0.55 | 5.07 | 0.33 | 6.18E+07 | ± 5.51E+06 |
| Alder_wet_3_15 | 74.9 | 78.4 | 630 | 96 | 302 | 141 | 2.77 | 34.0 | 12.3 | 0.55 | 5.07 | 0.33 | 1.02E+07 | ± 5.09E+05 |
| Alder_wet_3_25 | 80.3 | 52.1 | 791 | 91 | 350 | 171 | 2.71 | 39.0 | 14.4 | 0.65 | 5.09 | 0.33 | 6.68E+06 | ± 1.68E+05 |
| Coast_dry_1_05 | 44.2 | 19.2 | 422 | 54 | 182 | 91 | 1.59 | 21.7 | 13.6 | 0.59 | 3.85 | 1.32 | 4.29E+05 | ± 3.50E+04 |
| Coast_dry_1_15 | 62.4 | 30 | 470 | 58 | 218 | 100 | 1.17 | 15.1 | 12.9 | 0.59 | 4.05 | 1.32 | 4.20E+05 | ± 5.87E+04 |
| Coast_dry_1_25 | 91.0 | 74.9 | 1243 | 193 | 400 | 360 | 0.67 | 8.6 | 12.8 | 0.26 | 4.04 | 1.32 | 9.56E+05 | ± 2.59E+05 |
| Coast_dry_2_05 | 41.2 | 30.2 | 485 | 47 | 204 | 95 | 1.59 | 21.7 | 13.6 | 0.59 | 3.85 | 1.32 | 2.45E+05 | ± 4.45E+04 |
| Coast_dry_2_25 | 43.4 | 8.9 | 211 | 24 | 68 | 68 | 0.67 | 8.6 | 12.8 | 0.26 | 4.04 | 1.32 | 1.55E+05 | ± 7.36E+03 |
| Coast_dry_3_05 | 41.6 | 27.4 | 299 | 34 | 111 | 78 | 1.59 | 21.7 | 13.6 | 0.59 | 3.85 | 1.32 | 2.36E+05 | ± 2.73E+04 |
| Coast_dry_3_15 | 40.7 | 24.3 | 247 | 32 | 96 | 52 | 1.17 | 15.1 | 12.9 | 0.59 | 4.05 | 1.32 | 1.34E+05 | ± 3.57E+04 |
| Coast_dry_3_25 | 47.7 | 14.6 | 207 | 28 | 70 | 48 | 0.67 | 8.6 | 12.8 | 0.26 | 4.04 | 1.32 | 2.34E+05 | ± 5.44E+04 |
| Coast_wet_1_05 | 43.7 | 32.4 | 330 | 40 | 121 | 95 | 1.76 | 31.4 | 17.8 | 0.69 | 4.70 | 5.20 | 1.94E+06 | ± 1.79E+05 |
| Coast_wet_1_15 | 37.4 | 23.5 | 430 | 58 | 218 | 65 | 0.78 | 15.0 | 19.2 | 0.28 | 4.40 | 5.20 | 8.15E+05 | ± 3.27E+04 |
| Coast_wet_1_25 | 61.2 | 46.7 | 698 | 54 | 332 | 72 | 0.68 | 12.8 | 19.0 | 0.25 | 6.05 | 5.20 | 7.43E+05 | ± 5.32E+04 |
| Coast_wet_2_05 | 60.3 | 42.7 | 535 | 61 | 198 | 152 | 1.76 | 31.4 | 17.8 | 0.69 | 4.70 | 5.20 | 7.57E+06 | ± 1.06E+06 |
| Coast_wet_2_15 | 46.4 | 32.3 | 448 | 38 | 187 | 117 | 0.78 | 15.0 | 19.2 | 0.28 | 4.40 | 5.20 | 6.83E+05 | ± 3.09E+04 |
| Coast_wet_2_25 | 67.0 | 47.9 | 1050 | 73 | 393 | 393 | 0.68 | 12.8 | 19.0 | 0.25 | 6.05 | 5.20 | 9.91E+05 | ± 1.86E+05 |
| Coast_wet_3_05 | 54.5 | 54.1 | 550 | 44 | 223 | 114 | 1.76 | 31.4 | 17.8 | 0.69 | 4.70 | 5.20 | 2.20E+06 | ± 1.80E+05 |
| Coast_wet_3_15 | 45.7 | 36.7 | 418 | 63 | 223 | 78 | 0.78 | 15.0 | 19.2 | 0.28 | 4.40 | 5.20 | 5.95E+05 | ± 1.06E+05 |
| Coast_wet_3_25 | 68.1 | 42.8 | 763 | 77 | 368 | 172 | 0.68 | 12.8 | 19.0 | 0.25 | 6.05 | 5.20 | 5.97E+05 | ± 6.05E+04 |
| Perco_dry_1_05 | 62.4 | 95.9 | 825 | 88 | 477 | 173 | 3.41 | 37.3 | 10.9 | 0.49 | 4.98 | 0.11 | 1.40E+06 | ± 4.31E+05 |
| Perco_dry_1_15 | 73.6 | 110.5 | 1389 | 197 | 831 | 195 | 3.41 | 37.3 | 10.9 | 0.49 | 5.18 | 0.11 | 2.59E+06 | ± 2.48E+05 |
| Perco_dry_1_25 | 79.0 | 103.1 | 1024 | 175 | 606 | 229 | 3.07 | 43.6 | 14.2 | 0.50 | 5.20 | 0.11 | 7.22E+06 | ± 6.17E+05 |
| Perco_dry_2_05 | 71.4 | 102.3 | 938 | 91 | 486 | 97 | 3.41 | 37.3 | 10.9 | 0.49 | 4.98 | 0.11 | 1.18E+06 | ± 2.06E+05 |
| Perco_dry_2_15 | 75.4 | 117.9 | 944 | 113 | 530 | 124 | 3.41 | 37.3 | 10.9 | 0.49 | 5.18 | 0.11 | 2.08E+06 | ± 9.49E+04 |
| Perco_dry_2_25 | 78.4 | 73.7 | 810 | 97 | 419 | 105 | 3.07 | 43.6 | 14.2 | 0.50 | 5.20 | 0.11 | 2.11E+06 | ± 2.99E+06 |
| Perco_dry_3_05 | 71.6 | 125.6 | 817 | 86 | 441 | 118 | 3.41 | 37.3 | 10.9 | 0.49 | 4.98 | 0.11 | 2.33E+06 | ± 6.58E+05 |
| Perco_dry_3_15 | 76.6 | 181.4 | 1139 | 147 | 621 | 165 | 3.41 | 37.3 | 10.9 | 0.49 | 5.18 | 0.11 | 1.06E+07 | ± 7.81E+04 |
| Perco_dry_3_25 | 79.8 | 117.1 | 1143 | 132 | 647 | 156 | 3.07 | 43.6 | 14.2 | 0.50 | 5.20 | 0.11 | 9.50E+06 | ± 2.18E+06 |
| Perco_wet_1_05 | 83.5 | 195.4 | 1195 | 103 | 79 | 339 | 2.64 | 31.1 | 11.8 | 0.86 | 5.36 | 0.49 | 4.06E+07 | ± 2.06E+06 |
| Perco_wet_1_15 | 76.7 | 102.9 | 739 | 86 | 83 | 154 | 2.64 | 31.1 | 11.8 | 0.86 | 5.36 | 0.49 | 5.45E+07 | ± 6.35E+06 |
| Perco_wet_1_25 | 79.6 | 51.5 | 526 | 91 | 48 | 113 | 2.66 | 46.4 | 17.4 | 1.01 | 5.39 | 0.49 | 1.81E+07 | ± 3.02E+06 |
| Perco_wet_2_05 | 82.2 | 171.5 | 980 | 136 | 106 | 127 | 2.64 | 31.1 | 11.8 | 0.86 | 5.36 | 0.49 | 1.60E+07 | ± 1.33E+06 |
| Perco_wet_2_15 | 69.4 | 61.8 | 672 | 58 | 63 | 227 | 2.64 | 31.1 | 11.8 | 0.86 | 5.36 | 0.49 | 2.25E+07 | ± 1.22E+06 |
| Perco_wet_2_25 | 71.7 | 41.3 | 481 | 47 | 192 | 103 | 2.66 | 46.4 | 17.4 | 1.01 | 5.39 | 0.49 | 3.90E+07 | ± 6.14E+05 |
| Perco_wet_3_05 | 76.3 | 137.4 | 1682 | 104 | 782 | 351 | 2.64 | 31.1 | 11.8 | 0.86 | 5.36 | 0.49 | 4.92E+07 | ± 8.11E+06 |
| Perco_wet_3_15 | 71.0 | 81.3 | 1192 | 117 | 538 | 290 | 2.64 | 31.1 | 11.8 | 0.86 | 5.36 | 0.49 | 4.87E+07 | ± 1.27E+07 |
| Perco_wet_3_25 | 68.3 | 24.6 | 474 | 54 | 194 | 85 | 2.66 | 46.4 | 17.4 | 1.01 | 5.39 | 0.49 | 1.82E+07 | ± 1.83E+06 |
