## Supplementary material for "Rewetting of Three Drained Peatlands Drives Congruent Compositional Changes in Pro- and Eukaryotic Soil Microbiomes Through Environmental Filtering": Legends supplementory figures

Figure S1: Alpha diversity of fen microbiomes. a: prokaryotes: Shannon entropy of 16S rRNA amplicon sequencing ASV, bars show triplicates of cores, error bars indicate standard deviation between cores, b: eukaryotes: Shannon entropy of 18S rRNA amplicon sequencing ASV, bars show triplicates of cores, error bars indicate standard deviation between cores. A: Alder carr, C: coastal fen, P: percolation fen, D: drained, W: rewetted, depth in cm. Values are means of triplicate peat-cores. Kruskal-Wallis Test: *** significance level at p < 0.001; * significance level at p < 0.05, Pairwise T-test: significant difference between a and b.

Figure S2: Heatmap showing Spearman correlation between microbial community factors and soil parameters. Spearman correlation analysis between soil properties, alpha diversity, NMDS axes and abundances of functional groups and methanogenic orders. Color and size of points indicate coefficients, red indicate negative correlation, blue indicate positive correlation, the bigger the symbols the stronger the correlation. Water: gravimetric water content; DNA_DW: µg DNA per gram dry soil; DOM: dissolved organic matter; Bio: Biopolymers; hCOM: humic carbon organic matter; lCOM: low-molecular organic matter; Nanorg: anorganic nitrogen; D_NO3: dissolved nitrate; D_NO2:dissolved nitrite; D_NH4: dissolved Ammonia; N: total nitrogen; C: total carbon; C_N: carbon/nitrogen ratio; S: total sulfur; pH: pH from calcium chloride extracts;
Sal: salinity; mcrA_DNA: mcrA gene copies per ng DNA; mcrA_DW: mcrA gene copies per gram dry soil; alpha_16S/18S: alpha diversity of prokaryotic/eukaryotic community; MDS: multi-dimensional-scaling axis 1/2 for prokaryotes/eukaryotes; meth: relative abundance of methanogen; sulf_red: relative abundance of sulfate reducers; meth_troph: relative abundance of methanotrophs

Figure S3 Relative abundance of Acidobacteria in 16S rRNA gene amplicon datasets. Barplots show relative abundance. X-axes represent fen type: alder carr (A), coastal fen (C), percolation fen (P), hydrological state drained (D) and rewetted (W), depth in cm (05, 15, 25). Data are shown as mean values of triplicate soil cores.

Figure S4: Node features of co-occurrence network analysis. Factors of a) degree, b) closeness centrality and c) transitivity in drained and rewetted sites and d) degree, e) closeness centrality and f) transitivity in fen types Alder (A), Percolation (P) and coastal (C).
